# HDAC11 regulates type I interferon signaling through defatty-acylation of SHMT2

**DOI:** 10.1101/211706

**Authors:** Ji Cao, Lei Sun, Pornpun Aramsangtienchai, Nicole A. Spiegelman, Xiaoyu Zhang, Edward Seto, Hening Lin

## Abstract

The smallest histone deacetylase (HDAC) and the only class IV HDAC member, HDAC11, is reported to regulate immune activation and tumorigenesis, yet its physiological function is largely unknown. Here we identify HDAC11 as an efficient lysine defatty-acylase that is >10,000-fold more efficient than its deacetylase activity. Through proteomics studies, we identified SHMT2 as a defatty-acylation substrate of HDAC11. HDAC11-catalyzed defatty-acylation did not affect the enzymatic activity of SHMT2. Instead, it affects the ability of SHMT2 to regulate type I interferon receptor ubiquitination and internalization. Correspondingly, HDAC11 depletion increased type I interferon signaling in both cell culture and mice. This study is the first time a zinc-dependent HDAC is found to have an activity that is much more efficient than the corresponding deacetylase activity. The finding expands the physiological functions of HDAC11 and protein lysine fatty acylation, and opens up opportunities to develop HDAC11-specific inhibitors as therapeutics to modulate immune responses.

---

Histone deacetylases (HDACs) regulate many biological functions by removing acetyl groups from lysine residues on histones and non-histone proteins ^1,2^. There are 18 HDACs in mammals that are classified into four classes. Class III HDACs (known as sirtuins) are NAD^+^-dependent, while class I, II and IV HDACs are all zinc-dependent. Class I HDACs, which consists of HDAC1, 2, 3, and 8, are homologous to yeast Rpd3. Class II HDACs are homologous to yeast Hda1 and can be further divided into two subclasses: IIa (HDAC4, 5, 7, 9) and IIb (HDAC6, 10)^1,3^. Class IV HDAC, which is not highly homologous to either Rpd3 or Hda1 yeast enzymes, is comprised solely of HDAC11 ^4,5^.

HDAC11 is normally found in brain, testis, and immune cells such as antigen-presenting cells (APCs) ^5,6^. Upregulation of HDAC11 has been found in many cancer cells ^7,8^ and interleukin-13 (IL-13) treated B cells ^9^. It has been reported that HDAC11 negatively regulates IL-10 production in APCs by binding to the IL-10 promoter and promoting histone deacetylation, thus influencing immune activation versus tolerance ^10^. However, the deacetylase activity of HDAC11 reported in the literature is rather weak, and its ability to directly deacetylate histones has not been demonstrated. The lack of efficient deacetylase activity has prevented the identification of more substrate proteins that could help to further understand the biological function of HDAC11.

Recently, several HDACs have been shown to hydrolyze acyl lysine modifications that are distinct from acetyl lysine ^11–15^ In particular, SIRT5 can efficiently remove succinyl and malonyl groups ^16,17^, while SIRT6 can efficiently remove long chain fatty acyl groups in vitro and in vivo ^18^. These results prompted us to hypothesize that the zinc-dependent HDACs with weak or no deacetylase activities may also prefer to hydrolyze other acyl lysine modifications. Herein we identify an activity of HDAC11 that is >10,000 time more efficient than its deacetylase activity. This previously undiscovered, highly efficient, activity led us to identify a new lysine acylated protein and a regulatory pathway, which provides novel insights into the immune-modulatory function of HDAC11.

## Results

### HDAC11 is an efficient lysine defatty-acylase

In order to study the enzymatic activity of HDAC11, we first tested recombinant HDAC11 purified from HEK293T cells (Figure S1A) on H3K9 peptides bearing different acyl groups (Figure 1A). Using an HPLC-based assay, we found HDAC11 could efficiently remove long chain fatty-acyl groups (myristoyl, palmitoyl, and 3-hydroxydodecanoyl) from H3K9 peptides (Figure 1A and 1B). Surprisingly, HDAC11 failed to remove smaller acyl groups, including octanoyl and decanoyl, from H3K9 peptides (Figure 1A and 1B). Similarly, many other acyl groups tested, including lipoyl, succinyl, and glutaryl on H3K9 peptides were not HDAC11 substrates (Figure 1A). To further compare the deacetylation and demyristoylation activities of HDAC11, we also chose a few different peptide sequences. We detected demyristoylation activity of HDAC11 on H2BK12 and TNFα peptides, but did not detect any deacetylation activity on H2BK12 and α-tubulin peptides (Figure S1B). Since N-terminal glycine myristoylation is a well-known posttranslational modification ^19^, we also tested whether HDAC11 could catalyze glycine demyristoylation. No glycine demyristoylation activity was detected on p21-activated kinase 2 (PAK2) and Gα peptides (Figure S1C).

**Figure 1.**
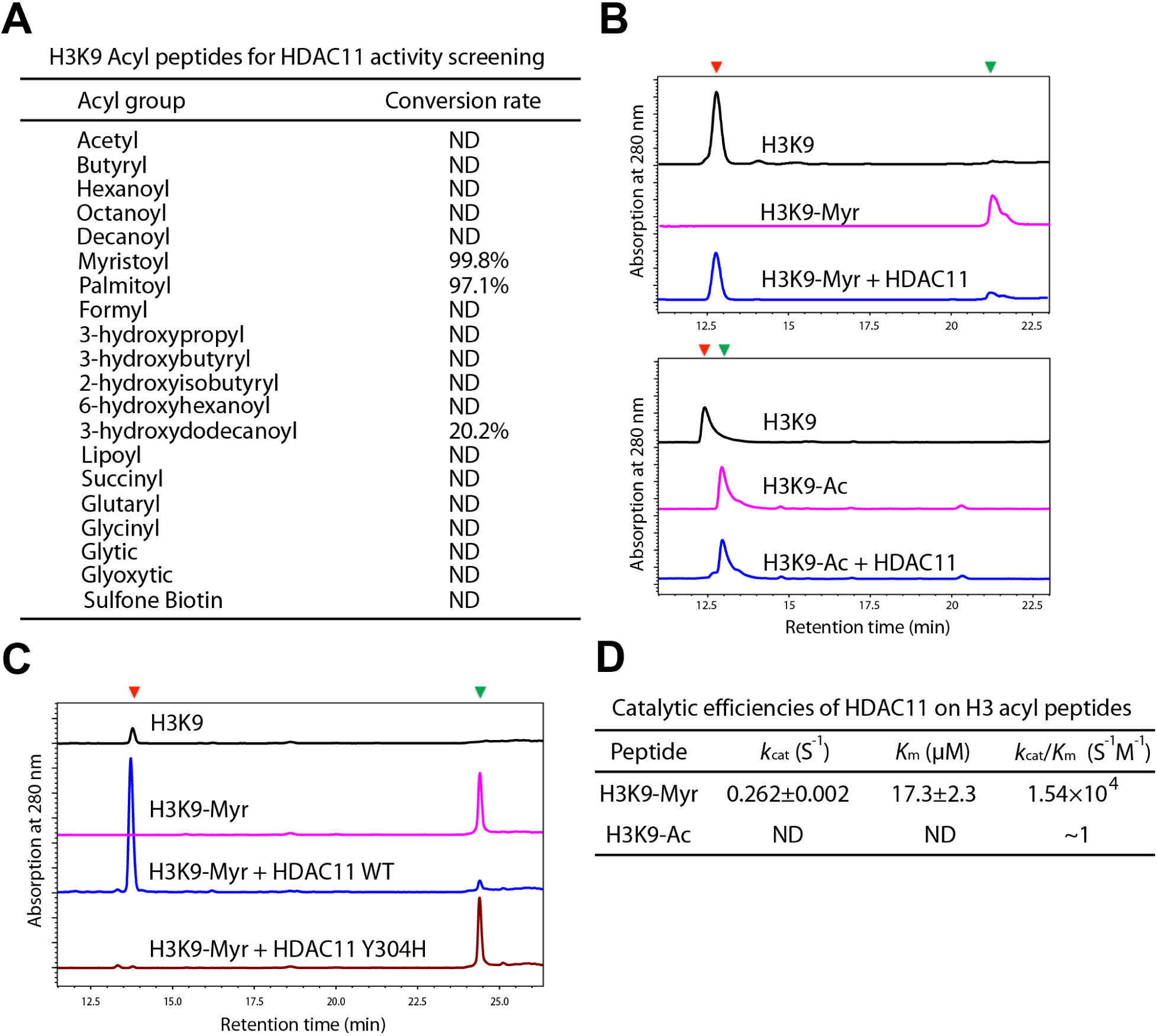
HDAC11 is an efficient lysine defatty-acylase. **A.** HDAC11 enzymatic activity on H3K9 peptides bearing different acyl groups. **B.** Representative data of HDAC11 enzymatic activity on myristoyl and acetyl H3K9 peptides. **C.** Representative data of wild type (WT) HDAC11 and its inactive mutant (Y304H) enzymatic activity on myristoyl H3K9 peptide. **D.** Kinetic parameters of HDAC11 on myristoyl and acetyl H3K9 peptides.

To rule out that the defatty-acylation activity was from a contaminating protein in the HDAC preparation, we further constructed and purified four catalytic mutants of HDAC11 that lack the general acid catalytic residue (Y304H) or zinc-binding residues (D181A, H183A, and D261A). Wild-type HDAC11, but not the catalytic mutants, could remove myristoyl group from the H3K9 peptide (Figure 1C and Figure S1D). Furthermore, we also detected the demyristoylation activity of recombinant HDAC11 purified from *Saccharomyces cerevisiae* (Figure S1E).

We next carried out kinetics studies using H3 K9 myristoyl peptide as the substrate. As shown in Figure 1D, the *K*_*m*_ value for the myristoyl H3K9 peptide was 17.3 μM and catalytic efficiency (*k*_*cat*_/*K*_*m*_ value) is 1.54×10^4^ M^−1^s^−1^. For deacetylation, we could not obtain the *k*_*cat*_/*K*_*m*_ value because no product was detected. But based on the detection limit of the HPLC assay, we estimated the upper limit of the *k*_*cat*_/*K*_*m*_ value to be about 1 M^−1^s^−1^. Thus, the lysine defatty-acylase activity of HDAC11 is >10,000-fold more efficient than its deacetylase activity in vitro.

### Proteomic approach identifies potential defatty-acylation substrates of HDAC11

To address whether the efficient defatty-acylation activity of HDAC11 is physiological relevant, we first tested the effect of HDAC11 knockdown (KD) on global lysine fatty-acylation level in MCF-7 cells, which had endogenous HDAC11 expression. Using a metabolic labeling method (Figure S2A) with the Alk14 probe (an alkyne-tagged fatty acid analogue) for protein myristoylation and palmitoylation ^18,20^, intracellular fatty-acylated proteins were labeled. The labeled proteins were then conjugated to a fluorescent tag (BODIPY-azide). After precipitating the proteins, hydroxylamine was used to remove cysteine acylation, and then the lysine acylated proteins were resolved by SDS-PAGE and visualized by in-gel fluorescence. As shown in Figure 2A, HDAC11 KD cells had increased global lysine fatty-acylation levels compared to control cells.

**Figure 2.**
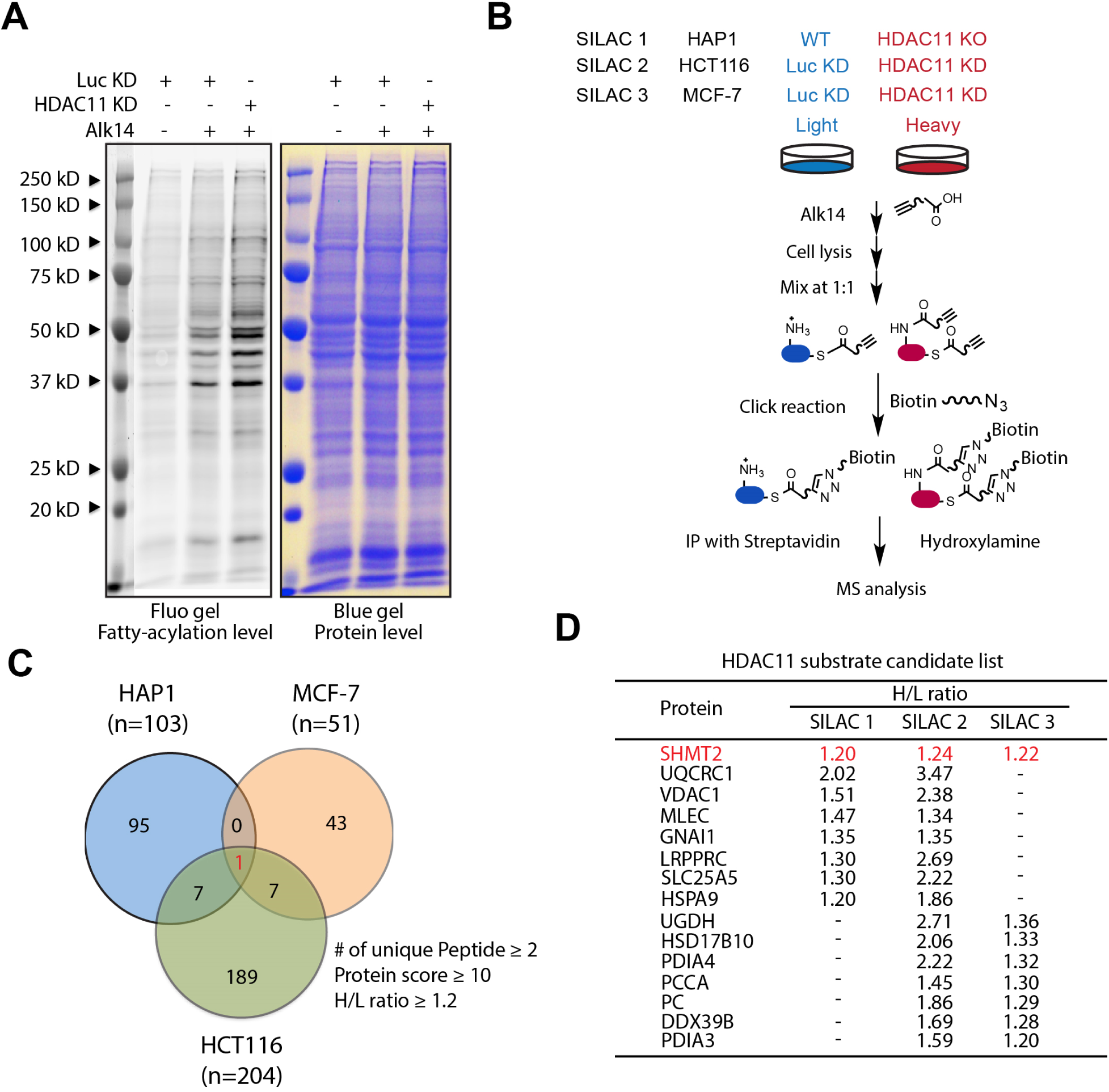
Proteomic approach identifies potential defatty-acylation substrates of HDAC11. **A.** The effect of HDAC11 knockdown (KD) on global lysine fatty-acylation level in MCF-7 cells using a metabolic labeling method with the Alk14 probe. **B.** Alk14 labeling and SILAC to identify proteins with increased Alk14 labeling in cells with HDAC11 KD or KO compared to control KD (Luc KD) or wild type cells. **C.** Potential substrates regulated by HDAC11 identified by SILAC in three pairs of cell lines (HAP1, HCT116, and MCF-7) with the following criteria: heavy/light (H/L) ratio .1.2 with at least two peptides and protein score ≥10. **D.** The 15 putative HDAC11 substrate proteins that were identified from at least two SILACs. The H/L ratios are listed.

We then set out to identify potential defatty-acylation substrates of HDAC11 using proteomics. As shown in Figure 2B, we combined Alk14 labeling and stable isotope labeling with amino acid in cell culture (SILAC) to identify proteins with increased Alk14 labeling in HDAC11 KD or knockout (KO) cells. The SILAC experiments were carried out using three pairs of cell lines: wild-type (WT) and HDAC11 knockout (KO) HAP1 cells (SILAC1), control and HDAC11 KD HCT116 cells (SILAC2), and control and HDAC11 KD MCF-7 cells (SILAC3). The HDAC11 KO or KD cells were cultured with heavy amino acids while the corresponding control cells were cultured with light amino acids. The cells were labeled with Alk14 and the alkyne-labeled proteins were conjugated to a biotin tag (biotin-azide) via click chemistry. After the modified proteins were pulled down with streptavidin and treated with hydroxylamine, they were analyzed by mass spectrometry. A protein with H/L ratio higher than 1 in SILAC could be a potential substrate of HDAC11. To remove false positives and simplify downstream validation studies, we further filtered our data using the following criteria: heavy/light (H/L) ratio ≥1.2 with at least two peptides and protein score ≥10. Based on these criteria, we identified 139, 212, and 98 proteins in SILAC1 (Table S1), SILAC2 (Table S2) and SILAC3 (Table S3), respectively. Among them, 31 proteins were identified from at least two SILACs (Figure 2C and Figure 2D). In particular, one protein, serine hydroxymethyltransferase 2 (SHMT2), was identified from all three SILACs with H/L ratios ≥1.2 (Figure 2D). We therefore focused further validation studies on SHMT2.

### SHMT2 is a defatty-acylation substrate of HDAC11 in cells

We first tested whether SHMT2 was a lysine fatty-acylated protein using Alk14 and fluorescent labeling (Figure S2B). Although some background fluorescent signal without Alk14 treatment was detected, Alk14 treatment increased the fluorescent signal of Flag-tagged SHMT2 (Flag-SHMT2, Figure 3A), suggesting that SHMT2 contained fatty-acylation. Moreover, we detected the fatty-acylation of endogenous SHMT2 in cells (Figure 3B) using Alk14 labeling, biotin-azide conjugation, and streptavidin pulldown, followed by immunoblotting against SHMT2. Therefore, our data suggested that SHMT2 is a fatty-acylated protein. Co-immunoprecipitation studies showed that overexpressed Flag-SHMT2 interacted with overexpressed HDAC11 with an HA tag in HEK 293T cells (Figure S2C).

**Figure 3.**
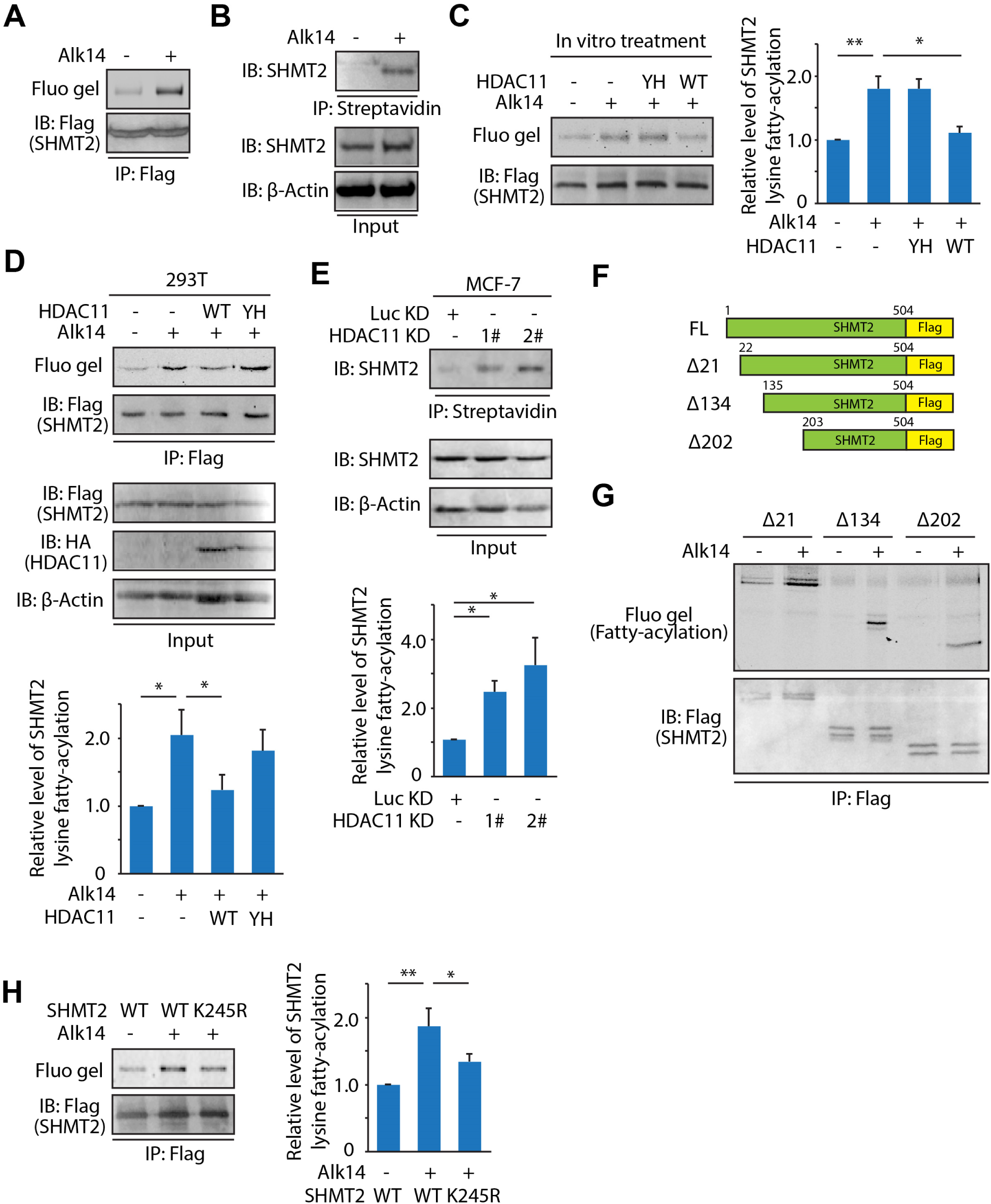
SHMT2 is a defatty-acylation substrate of HDAC11 in cells. **A.** In-gel fluorescence showing SHMT2 is a lysine fatty-acylated protein using Alk14 labeling and fluorescence conjugation through click chemistry. **B.** Fatty-acylation of endogenous SHMT2 in HEK293T cells was detected using Alk14 labeling, biotin-azide conjugation, and streptavidin pulldown, followed by immunoblotting against SHMT2. **C.** SHMT2 Alk14 labeling after *in vitro* treatment with wild-type (WT) or the Y304H catalytic mutant (YH) of HDAC11. Quantification of the relative levels of lysine fatty-acylation was shown in the right panel. **D.** In-gel fluorescence showing lysine fatty-acylation of SHMT2 was regulated by WT but not the Y304H catalytic mutant of HDAC11 in HEK293T cells co-overexpressing SHMT2 and HDAC11. Quantification of the relative levels of lysine fatty-acylation was shown in the lower panel. **E.** The fatty-acylation of endogenous SHMT2 was increased after knocking down HDAC11 in MCF-7 cells. Quantification of the relative level of lysine fatty-acylation was shown in the lower panel. **F.** Schematic representation of Flag-tagged WT and truncated SHMT2 constructs. **G.** In-gel fluorescence showing fatty-acylation was localized to residue 203-504 of SHMT2. **H.** In-gel fluorescence gel showing K245 is the major fatty-acylation site of SHMT2. Quantification of the relative level of lysine fatty-acylation is shown in the right panel. In all panels, values with error bars indicate mean ± sd of three replicates.

To determine whether SHMT2 is a direct substrate of HDAC11, we first examined whether recombinant HDAC11 could remove the lysine fatty-acylation on SHMT2 in vitro. Flag-SHMT2 was overexpressed in 293T cells and labeled with Alk14. Immunoprecipitated SHMT2 was incubated with wild-type (WT) HDAC11 or the Y304H catalytic mutant (YH) of HDAC11. Then a fluorescent tag (BODIPY-azide) was conjugated using click chemistry and the fatty acylation level on SHMT2 was detected using in-gel fluorescence. As shown in Figure 3C, wild-type, but not mutant, HDAC11 significantly decreased the fluorescent signal on SHMT2, suggesting that HDAC11 can directly remove the fatty acyl groups on SHMT2.

In HEK 293T cells, co-overexpression of SHMT2 and WT HDAC11 decreased the fatty-acylation level on SHMT2, compared to cells without HDAC11 overexpression or cells overexpressing the Y304H catalytic mutant (Figure 3D). Thus, the enzymatic activity of HDAC11 is required for controlling SHMT2 fatty acylation in cells. Additionally, in MCF-7 cells, HDAC11 KD significantly increased endogenous SHMT2 fatty-acylation level compared to control KD (Figure 3E). Similar results were also observed in A549 and HCT116 cell lines (Figure S3A). Collectively, our *in vitro* and cellular data support the hypothesis that HDAC11 regulates the fatty-acylation level of SHMT2.

We next examined which lysine residue of SHMT2 was the fatty-acylation site. We first constructed a series of truncated forms of SHMT2 (Δ21, Δ134 and Δ202) (Figure 3F) to narrow down the modification region. As shown in Figure 3G, all three truncated forms still had lysine fatty-acylation, suggesting the modified lysine is localized between residues 203 and 504 of SHMT2, which contained 16 lysine residues. Then, we mutated each of the 16 lysine (K) to arginine (R). Among them, 14 mutants were successfully expressed in HEK 293T cells (Figure S3B) while the other two could not be expressed. Our results showed that only the K245R mutant significantly decreased the fatty acylation level, suggesting K245 is the major acylation site (Figure 3H and Figure S3B). Interestingly, this lysine residue is not conserved in another isoform of SHMT in mammals, SHMT1. SHMT1 did not contain fatty acylation (Figure S3C and Figure S3D), which further supported that K245 is the major lysine fatty acylation site in SHMT2.

### HDAC11 defatty-acylation of SHMT2 regulates IFNαRI internalization and stability

We next investigated the biological function of SHMT2 lysine fatty acylation. SHMT2, a metabolic enzyme involved in one-carbon metabolism, catalyzes the reversible interconversion of serine and tetrahydrofolate to glycine and methylenetetrahydrofolate ^21,22^. Thus, we first tested whether fatty-acylation might affect the enzymatic activity of SHMT2. As shown in Figure 4A, the enzymatic activity of SHMT2 K245R mutant was not significantly different from that of wild type SHMT2. Similarly, the enzymatic activity of SHMT2 was not significantly changed when co-expressed with either wild type or catalytic mutant of HDAC11 in 293T cells (Figure 4B). These observations suggested fatty-acylation did not affect SHMT2 enzymatic activity. In line with these data, we also did not detect any difference in the oligomeric states of K245R mutant and wild type SHMT2 (Figure S4A), which forms an active tetramer ^23^.

**Figure 4.**
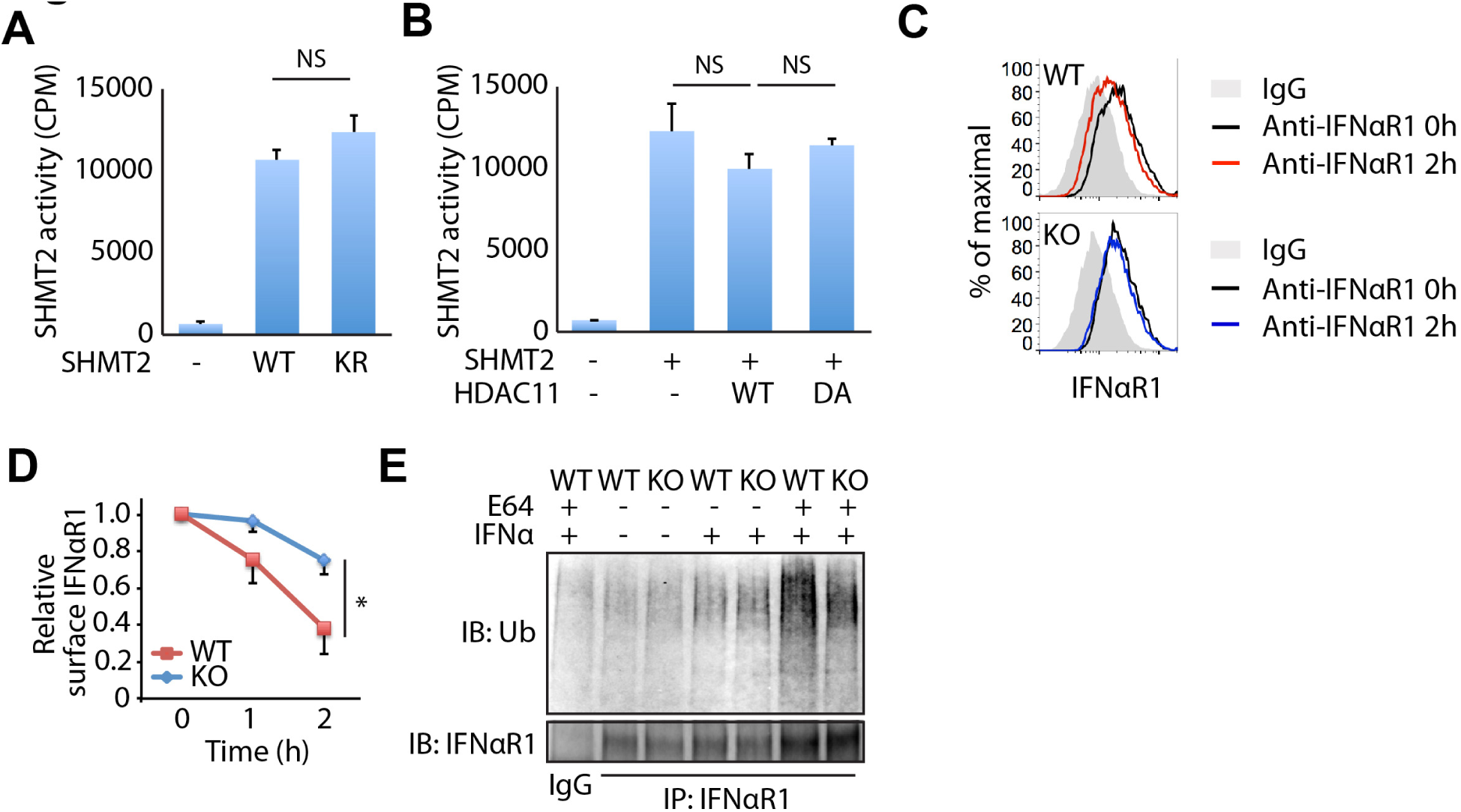
HDAC11 defatty-acylation of SHMT2 regulates IFNαRI internalization and stability. **A.** The enzymatic activity of WT and the K245R mutant (KR) of SHMT2 purified from 293T cells. **B.** The enzymatic activity of SHMT2 co-overexpressed with WT and the Y304H catalytic mutant of HDAC11 in 293T cells. C, D. The internalization rate of IFNαRI upon IFNα treatment in WT and HDAC11 KO HAP1 cells by monitoring cell surface IFNαRI level using flow cytometry. **C.** The representative data of cell surface IFNaR1 levels before and after 2h IFNα treatment in WT and HDAC11 KO HAP1 cells. **D.** The relative surface IFNaR1 level upon IFNα treatment in WT and HDAC11 KO HAP1 cells with indicated time points. **E.** The ubiquitination level of IFNaR1 after treating wild-type and HDAC11 KO HAP1 cells with IFNα. The western blotting result is the representative image from three independent experiments. Statistical analysis: values with error bars indicate mean ± sd of three replicates and NS indicates no statistical difference.

SHMT2 is mainly localized in the mitochondria and the subcellular localization of wild type SHMT2 and the K245R mutant in wild type or HDAC11 KO HAP1 cells was not significantly different (Figure S4B and S4C). However, we did detect a small amount of SHMT2 that was localized in the cytosol (Figure S4C), which was consistent with previous reports ^21,24^. Given that the natural cytosolic isoform of SHMT2 (also known as SHMT2α, referred to as Δ21 mutant here) also had lysine fatty-acylation (Figure 3E and S4D), we concluded that lysine fatty acylation mainly occurred to the cytosolic SHMT2α. Cytosolic SHMT2 was known to direct the BRCC36 containing complex (BRISC, a complex with deubiquitination activity) to deubiquitinate type I interferon receptor chain 1 (IFNαR1), and thus decrease the internalization and increase the stability of IFNαR1 ^24^ We thus tested whether HDAC11 affects the internalization and stability of IFNαR1. As shown in Figure 4C and 4D, cell surface IFNαR1 dramatically decreased due to internalization upon IFNα treatment in wild type cells, but not in HDAC11 KD cells. Consistent with this, HDAC11 knockout dramatically decreased the ubiquitination level of IFNαR1 compared to wild type cells (Figure 4E). Therefore, our data suggested that HDAC11 defatty-acylation of SHMT2α promoted IFNαR1 ubiquitination and internalization.

### HDAC11 regulates type I interferon signaling

IFNαR1 is important for type I interferon signaling, which controls the activation of many genes that are important for immune response ^25^. The protein stability of IFNαR1 would affect the activation of downstream genes in type I interferon treated cells ^24,26^. Thus, HDAC11 may regulate type I interferon signaling via defatty-acylation of SHMT2. To test this, we measured expression of two classic IFNα-driven genes, *ISG15* and *PKR ^25^.* As shown in Figure 5A and 5B, *ISG15* and *PKR* mRNA levels were increased in HDAC11 knockout cells more than those in WT HAP1 cells. Similarly, *ISG15* and *PKR* mRNA levels were higher in HDAC11 KD MCF-7 cells than in control KD MCF-7 cells. As expected, re-expressing mouse HDAC11 in HDAC11 KO HAP1 cells diminished the increase of *ISG15* and *PKR* expression (Figure S5A). Furthermore, knockdown of either SHMT2 or KIAA0157, key components of BRSIC ^24^, diminished the increase of *ISG15* and *PKR* mRNA level in HDAC11 knockout HAP1 cells (Figure 5C and S5B), further supporting the idea that HDAC11 antagonizes type I interferon signaling via defatty-acylation of SHMT2α, decreasing the recruitment of BRISC, and increasing the ubiquitination of IFNαRl.

**Figure 5.**
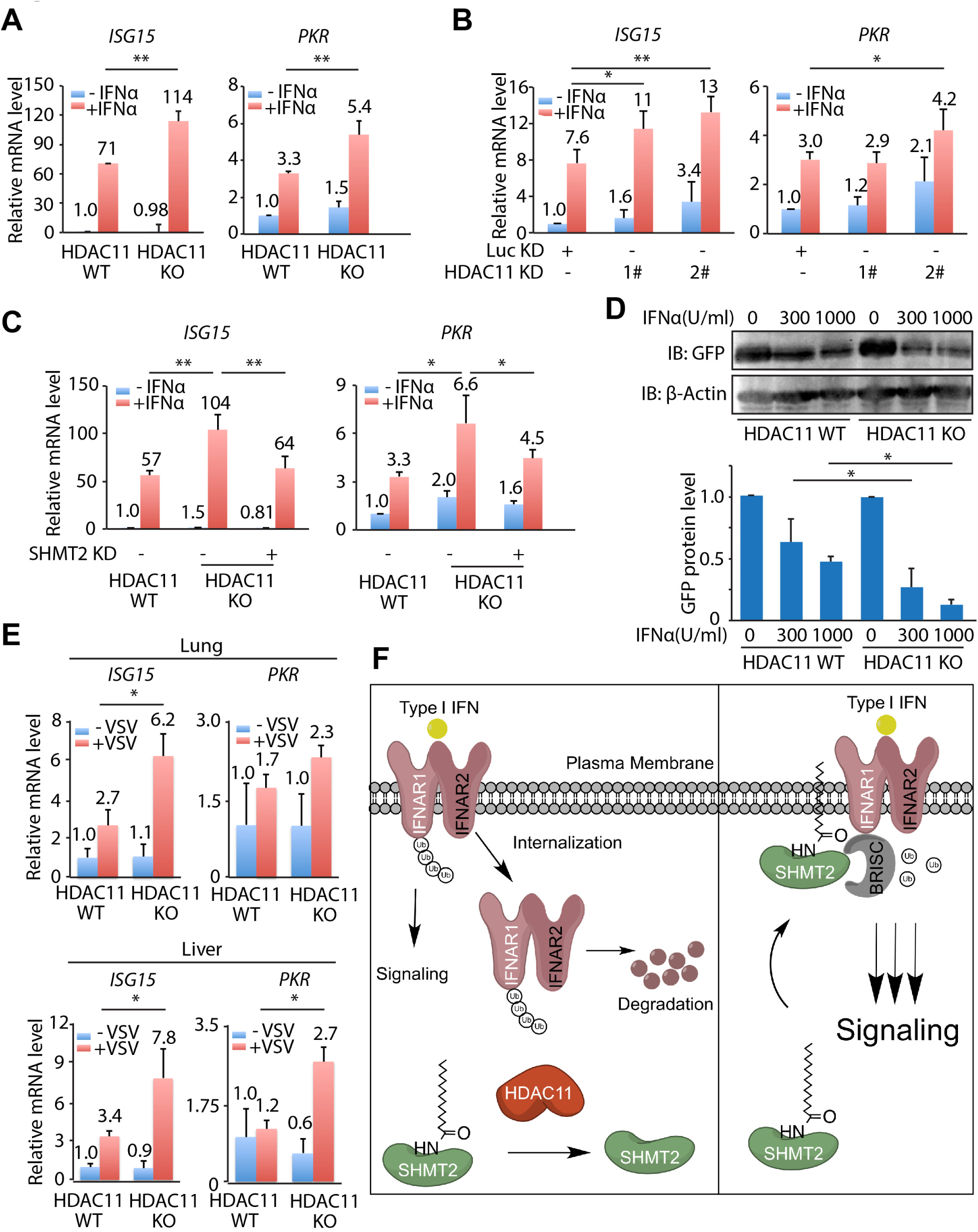
HDAC11 regulates type I interferon signaling. **A.** The relative mRNA levels of IFNαRI downstream genes, *ISG15* and *PKR,* in IFNα treated WT and HDAC11 KO HAP1 cells were quantified by RT-PCR and normalized to GAPDH expression. B. The relative mRNA levels of *ISG15* and *PKR* in IFNα treated control (Luc KD) and HDAC11 KD MCF-7 cells were quantified by RT-PCR and normalized to GAPDH expression. C. The effect of SHMT2 KD on the relative mRNA level of *ISG15* and *PKR* in HDAC11 KO HAP1 cells. D. HDAC11 KO cells had reduced GFP expression compared to WT HAP1 cells when treated with IFNα and infected with lentivirus carrying a pCCL vector encoding GFP. Quantification of the GFP protein levels is shown in the lower panel. E. WT and HDAC11 KO mice were injected with VSV for 2 days and the lung and liver were harvested for mRNA analysis. The relative mRNA levels of IFNαR1 downstream genes, *ISG15* and *PKR,* were quantified by RT-PCR and normalized to GAPDH expression. Values with error bars indicate mean ± sd of three samples.

Type I interferon is released by mammalian cells in response to viral infections and helps cells to heighten their anti-viral defense ^25^. Therefore, we further investigated whether HDAC11 could affect the viral defense of cells. We treated WT and HDAC11 knockout HAP1 cells with IFNα before infecting cells with lentiviruses carrying the GFP gene in a pCCL vector. The protein expression level of GFP would be reduced if there was increased anti-viral defense due to increased IFNαRl signaling in HDAC11 knockdown cells. Consistent with our hypothesis, HDAC11 knockout cells had reduced GFP protein levels compared to wild type HAP1 cells (Figure 5D).

To validate this phenotype *in vivo,* we further examined the IFNαRI signaling using transgenic mice model infected with vesicular stomatitis virus (VSV). The *ISG15* and *PKR* mRNA levels in the lung and liver was measured after infecting with VSV for 2 days. As shown in Figure 5E, VSV viral infection upregulated the *ISG15* and *PKR* mRNA level as expected. In line with the role of HDAC11 in regulating IFNαR1 signaling in cells, we observed a significant increase of *ISG15* and *PKR* mRNA level in lung and liver tissues from VSV-infected HDAC11 knockout mice comparing to those from wild-type animals (Figure 5E), consistent with the model that HDAC11 could antagonize type I interferon signaling (Figure 5F).

## Discussion

In this study, we have identified HDAC11 as an efficient lysine defatty-acylase with catalytic efficiency >10,000 fold better than its deacetylase activity. This is the first time a zinc-dependent HDAC is found to have an activity that is much more efficient than its corresponding deacetylase activity. HDAC8 was recently reported to have defatty-acylase activity, but the catalytic efficiency is only 2-3 fold better than that of deacetylation ^12^. The studies with HDAC11 suggest that the zinc-dependent HDACs are similar to the sirtuin family of HDACs, where several members preferentially recognize acyl lysine modifications other than acetyl lysine, while some can remove multiple modifications with similar efficiencies ^11,13^. Among the zinc-dependent HDACs, class IIa HDACs lack detectable deacetylase activity ^1^. The finding that HDAC11 has a novel efficient activity uncovers the possibility that class IIa HDACs may also be able to remove currently unknown acyl lysine modifications.

Our study identified a novel posttranslational mechanism that regulates a multifunctional protein, SHMT2 (Figure 5F). SHMT2 is a mitochondrial enzyme involved in one carbon metabolism ^21,22^. It has been reported that a shorter isoform, SHMT2α, which lacks the N-terminal mitochondrial localization sequence, is present in the cytosol and nucleus ^21^. The metabolic function of SHMT2 is thought to be critical for cancer cells and knockdown of SHMT2 inhibits the proliferation of several human cancer cells ^27^. Recently, SHMT2α was also reported to recruit the BRISC complex to IFNαR1 to promote the deubiquitination of IFNαRI ^24^. Thus, SHMT2 appears to have multiple functions. Our study presented here indicate that SHMT2α is regulated by lysine fatty acylation, which has only been known to occur on a few proteins in mammalian cells.^28–31^ Our data suggest that lysine fatty acylation does not alter the enzymatic activity of SHMT2α, but instead promotes the recruitment of the BRISC complex to IFNαR1 (Figure 5F).. HDAC11 can remove the lysine fatty acylation on SHMT2α, and thus increase the ubiquitination of IFNαR1 and downregulate interferon signaling (Figure 5F).

Our results revealed a new immune-regulatory function of HDAC11 by connecting it to the interferon signaling pathway. It has been reported that HDAC11 overexpression suppresses LPS stimulated IL-10 transcription while HDAC11 knockdown increases LPS-stimulated IL-10 transcription ^10^. This has been explained by a model where HDAC11 binds to IL-10 promoter and deacetylates histones to suppress IL-10 transcription ^10^. In this model, HDAC11 functions epigenetically at the end of the signaling pathway. In contrast, the HDAC11-IFNαR1 (Figure 5F) regulation that we discovered here occurs at the beginning of the signaling pathway. Our model is based on the more efficient defatty-acylase activity of HDAC11, while the previous model is based on the deacetylase activity of HDAC11. The two models are not mutually exclusive, or contradictory, since it is possible that HDAC11’s deacetylase activity could be increased upon chromatin binding which allows for deacetylation of histones, while it functions efficiently as defatty-acylase in the cytosol. A similar mechanistic function has been reported for SIRT6, a member of the NAD^+^-dependent HDAC ^31,32^.

We demonstrated here that the regulation of the interferon signaling pathway is physiologically relevant for the antiviral response in human cell lines and in mice. Mice lacking HDAC11 can increase interferon signaling, which may subsequently more effectively clear viral infections. This finding significantly expands the physiological function of HDAC11 and opens up opportunities to develop HDAC11-specific inhibitors as therapeutics to treat disease where increased type I interferon signaling is beneficial, such as viral infection,^33^ multiple sclerosis,^34^ or cancer ^35^.

## Materials and Methods

### Reagents

Anti-Flag affinity gel (cat log # A2220) and anti-Flag antibody conjugated with horseradish peroxidase (A8592) were purchased from Sigma-Aldrich. Antibodies against HDAC11 (sc-390737), β-actin (sc-4777), SHMT2 (sc-390641), GFP (sc-8334), and HA (sc-516102) were purchased from Santa Cruz Biotechnology. IFNαRI (LS-C290931) antibody was purchased from LifeSpan BioSciences. HSP60 (611563) antibody was purchased from BD Biosciences. Triple Flag peptide, protease inhibitor cocktail, [^13^C_6_,^15^N_2_]-L-lysine and [^13^C_6_,^15^N_4_]-L-arginine were purchased from Sigma-Aldrich. Sequencing-grade modified trypsin and FuGene 6 Transfection Reagent were purchased from Promega. Streptavidin agarose beads, ECL plus western blotting detection reagent and universal nuclease for cell lysis were purchased from Thermo Scientific Pierce. Sep-Pak C18 cartridges were purchased from Waters. Human interferon alpha 2a (11101-2) was purchased from PBL Assay Science.

### Cell lines and plasmids construction

The CRISPR-cas9-based HAP1 HDAC11 KO and wild type HAP1 cell lines were purchased from Horizon (Cambridge, UK). HAP1 cells were cultured in IMDM medium supplemented with 10% FBS in a humidified atmosphere of 5% CO_2_ at 37°C. The MCF-7, HCT116, A549 and HEK293T cells were purchased from ATCC and cultured in DMEM or RPMI-1640 medium supplemented with 10% FBS (Invitrogen) in a humidified atmosphere with 5% CO_2_ at 37°C.

To generate stable overexpression cells, lentivirus was generated by co-transfection of pCDH containing the desired gene, pCMV-dR8.2, and pMD2.G plasmids into HEK293T cells. The cell medium was collected 48 h after transfection and used to infect cells of interest. After 72h, infected cells were further treated with 1.5 mg/mL puromycin to select for stable overexpression cells. Empty pCDH vector was used as negative controls.

The full-length coding sequences for human HDAC11 and SHMT2 were amplified from MCF-7 cDNA library, and mouse HDAC11 was amplified from MEF cDNA library using Platinum Pfx DNA polymerase (ThermoFisher). The genes were subsequently sub-cloned into the pCMV-tag 4A, pcDNA3.0-3’HA and/or pCDH plasmid (CD510B-1) with the following primers:

pCMV-HDAC11 forward:AGTCAG<u>GGATCC</u>ATGCTACACACAACCCAGCTGTACCAG
pCMV-HDAC11 reverse:AGTCAG<u>GTCGAC</u>GGGCACTGCAGGGGGAAGCAGCGGT
pcDHA3.0-HA-HDAC11 forward: AGTCAG<u>GAATTC</u>ATGCTACACACAACCCAGCTGTACCAG
pcDHA3.0-HA-HDAC11 reverse: AGTCAG<u>GTCGAC</u>GGGCACTGCAGGGGGAAGCAGCGGT
pCMV-SHMT2 forward:AGTCAG<u>GAATTC</u>ATGCTGTACTTCTCTTTGTTTTGGGCG
pCMV-SHMT2 reverse: AGTCAG<u>CTCGAG</u>ATGCTCATCAAAACCAGGCATGGGGAAG
pCDH-SHMT2 forward:AGTCAG<u>TCTAGA</u>ATGCTGTACTTCTCTTTGTTTTGGGCG
pCDH-SHMT2 reverse:AGTCAG<u>GAATTC</u>CTACTTATCGTCGTCATCCTTGTAATC
pCDH-mHDAC11 forward: AGTCAG<u>GAATTC</u>ATGCCTCACGCAACACAGCTGTACCA
pCDH-mHDAC11 reverse: CAG<u>GGATCC</u>TTAAGCGTAATCTGGAACATCGTATGGGTAAGGCACAGCACAGGAAAGCAG
pCMV-SHMT2Δ21 forward: AGTCAG<u>GAATTC</u>ATGGCCATTCGGGCTCAGCACA
pCMV-SHMT2Δ134 forward: AGTCAG<u>GAATTC</u>ATGTGGGGAGTCAATGTCCAGCCCTACT
pCMV-SHMT2Δ202 forward: AGTCAG<u>GAATTC</u>ATGGGCCTCATTGACTACAACCAGCTGGCA
pCMV-SHMT2truncation reverse:AGTCAG<u>CTCGAG</u>ATGCTCATCAAAACCAGGCATGGGGAAG

The mutants of HDAC11 and SHMT2 were made by Phusion High-Fidelity DNA Polymerase Kit (New England Biolabs) with the following primers:

HDAC11 D181A forward:ACCATCATT GATCTTGCT GCCCAT CAGGGCAAT
HDAC11 D181A reverse:AAGATCAAT GATGGTAGCCCTGGAGATGCCCTC
HDAC11 H183A forward:ATTGATCTTGATGCCGCTCAGGGCAATGGGCAT
HDAC11 H183A reverse:GGCATCAAGATCAATGATGGTAGCCCTGGAGAT
HDAC11 D261A forward:TACAATGCAGGCACCGCCATCCTCGAGGGGGAC
HDAC11 D261A reverse:GGTGCCTGCATTGTATACCACCACGTCGGGCAG
HDAC11 Y304H forward: GTGACCTCAGGCGGGCACCAGAAGCGCACAGCC
HDAC11 Y304H reverse: CCCGCCTGAGGTCACCATAAGGATGGGCACCCG
SHMT2 K245R forward:GTGTGTGATGAAGTCCGAGCACACCTGCTGGCA
SHMT2 K245R reverse:GACTTCATCACACACCTCTCTCATGCGGGCGTA
SHMT2 K262R forward:GGCCTGGTGGCTGCCCGGGTGATTCCCTCGCCT
SHMT2 K262R reverse:GGCAGCCACCAGGCCACTGATGTGGGCCATGTC
SHMT2 K269R forward:ATTCCCTCGCCTTTCCGGCACGCGGACATCGTC
SHMT2 K269R reverse:GAAAGGCGAGGGAATCACCTTGGCAGCCACCAG
SHMT2 K280R forward:ACCACCACTACTCACCGGACTCTTCGAGGGGCC
SHMT2 K280R reverse:GTGAGTAGTGGTGGTGACGATGTCCGCGTGCTT
SHMT2 K294R forward:CTCATCTTCTACCGGCGAGGGGTGAAGGCTGTG
SHMT2 K294R reverse:CCGGTAGAAGATGAGCCCTGACCTGGCCCCTCG
SHMT2 K297R forward:TACCGGAAAGGGGTGCGGGCTGTGGACCCCAAG
SHMT2 K297R reverse:CACCCCTTTCCGGTAGAAGATGAGCCCTGACCT
SHMT2 K302R forward:AAGGCTGTGGACCCCCGGACTGGCCGGGAGATC
SHMT2 K302R reverse:GGGGTCCACAGCCTTCACCCCTTTCCGGTAGAA
SHMT2 K340R forward:GTAGCTGTGGCCCTACGGCAGGCCTGCACCCCC
SHMT2 K340R reverse:TAGGGCCACAGCTACTGCAGCAATGGCATGATT
SHMT2 K356R forward:TCCCTGCAGGTTCTGCGGAATGCTCGGGCCATG
SHMT2 K356R reverse:CAGAACCTGCAGGGAGTACTCCCGGAACATGGG
SHMT2 K389R forward:GTGGACCTGCGGCCCCGGGGCCTGGATGGAGCT
SHMT2 K389R reverse:GGGCCGCAGGTCCACCAGCACCAGGTGGTTGTC
SHMT2 K409R forward:TCCATCACTGCCAACCGGAACACCTGTCCTGGA
SHMT2 K409R reverse:GTTGGCAGTGATGGATACAAGCTCTAGCACCCG
SHMT2 K459R forward:ATTGGCTTAGAGGTGCGGAGCAAGACTGCCAAG
SHMT2 K459R reverse:CACCTCTAAGCCAATGTTGACCCCTTCATCTAT
SHMT2 K461R forward:TTAGAGGTGAAGAGCCGGACTGCCAAGCTCCAG
SHMT2 K461R reverse:GCTCTTCACCTCTAAGCCAATGTTGACCCCTTC
SHMT2 K464R forward:AAGAGCAAGACTGCCCGGCTCCAGGATTTCAAA
SHMT2 K464R reverse:GGCAGTCTTGCTCTTCACCTCTAAGCCAATGTT
SHMT2 K469R forward:AAGCTCCAGGATTTCCGATCCTTCCTGCTTAAG
SHMT2 K469R reverse:GAAATCCTGGAGCTTGGCAGTCTTGCTCTTCAC
SHMT2 K474R forward:AAATCCTTCCTGCTTCGGGACTCAGAAACAAGT
SHMT2 K474R reverse:AAGCAGGAAGGATTTGAAATCCTGGAGCTTGGC

### Generation of knockdown cell lines by shRNA

All shRNA lentivirus plasmids were purchased from Sigma. shRNA targeting luciferase was used as negative control. Lentivirus was generated by co-transfection of shRNA lentivirus plasmid, pCMV-dR8.2, and pMD2.G into HEK293T cells. The medium was collected 48 h after transfection and used to infect cells. After 72h, cells were used for indicated experiments. For stable knockdown, cells were further treated with 1.5 μg/mL puromycin to select for stable knockdown cells.

Luciferase shRNA:

CCGGCGCTGAGTACTTCGAAATGTCCTCGAGGACATTTCGAAGTACTCAGCGTTTTTG

HDAC11 shRNA 1#:

CCGGGTTTCTGTTTGAGCGTGTGGACTCGAGTCCACACGCTCAAACAGAAACTTTTTG

HDAC11 shRNA 2#:

CCGGGCGCTATCTTAATGAGCTCAACTCGAGTTGAGCTCATTAAGATAGCGCTTTTTG

SHMT2 shRNA:

CCGGACAAGTACTCGGAGGGTTATCCTCGAGGATAACCCTCCGAGTACTTGTTTTTTG

KIAA0157 shRNA:

CCGGGGGCGATTTATCAGGTTTATACTCGAGTATAAACCTGATAAATCGCCCTTTTTG

### Expression and purification of human HDAC11 from HEK293T cells

HDAC11 plasmids (WT, D181A, H183A, D261A, and Y304H) were transfected into HEK293T cells using FuGene 6 transfection reagent following the manufacturer’s protocol (Promega). The pCMV-Tag 4a empty vector was used as the negative control. The cells were collected by centrifugation at 1000 g for 5 min and then lysed in NP-40 lysis buffer (25 mM Tris-HCl, pH 7.8, 150 mM NaCl, 10% glycerol and 1% NP-40) with protease inhibitor cocktail (1:100 dilution) at 4 °C for 30 min. After centrifugation at 17,000 g for 30 min, the supernatant was collected and incubated with 20 μL of anti-Flag affinity gel at 4 °C for 2 h. The affinity gel was washed three times with washing buffer (25 mM Tris-HCl, pH 7.8, 150 mM NaCl, 0.2% NP-40) and then eluted with 300 μM of triple FLAG peptide (dissolved in 25 mM Tris-HCl, pH 7.4, 150 mM NaCl, and 10% glycerol). Eluted HDAC11 was used for the deacylation assay.

### *In Vitro* HDAC11 Activity and Kinetics Assay

HDAC11 *in vitro* activity was detected using an HPLC assay. Different acyl peptides were synthesized as previously reported ^12,36^. Each enzymatic reaction consisted of 25 μM of an acyl peptide in the reaction buffer (25 mM Tris-HCl pH 7.8, 150 mM NaCl) with a final volume of 50 μL. To initiate the reaction, 120 nM of HDAC11 was added into the reaction and incubated at 37 °C for 2 h. The reactions were quenched with 50 μL of acetonitrile and centrifuged at 17 000 g for 5 min. The supernatant was analyzed by reverse phase HPLC with a Kinetex 5U XB C18 column (100Å, 150 mm × 4.60 mm, Phenomenex) monitoring at 280 nm. The mobile phase A was water with 0.1% (v/v) trifluoroacetic acid (TFA); the mobile phase B was acetonitrile with 0.1% (v/v) TFA. The gradient used was 10 to 100% mobile phase B over 30 min with the flow rate of 0.5 mL/min. The product formation was verified by LC-MS (LCQ Fleet, Thermo Scientific).

The kinetics parameters of HDAC11 on the myristoyl peptide were determined using the above reaction conditions and by varying the concentrations of the myrisotyl H3K9 peptide between 0 and 200 μM. Each reaction was incubated with 50 nM of HDAC11 at 37 °C for 1 h. Each reaction was performed in duplicate to ensure reproducibility. After separation by HPLC, the product and remaining substrate peaks were quantified and converted to initial rates. The plots between substrate concentrations and the initial rates were fitted to the Michaelis-Menten equation using GraphPad Prism to obtain *k*_*cat*_ and *K*_*m*_ values.

### Detection of lysine fatty acylation on SHMT2 by in-gel fluorescence

The SHMT2 plasmid and/or different forms of HDAC11 plasmids (WT, YH, DA) were transfected into HEK293T cells using FuGene 6 transfection reagent. After 24 h, the cells were treated with 50 μM Alk14 for another 6 h. The cells were collected by centrifugation at 1000 g for 5 min and then lysed in NP-40 lysis buffer (25 mM Tris-HCl, pH 7.4, 150 mM NaCl, 10% glycerol, and 1*%* NP-40) with protease inhibitor cocktail (1:100 dilution) at 4 °C for 30 min. After centrifuging at 15,000 g for 15 min, the supernatant was collected and incubated with 10 μL of anti-Flag affinity gel at 4 °C for 1.5 h. The affinity gel was washed three times with washing buffer (25 mM Tris-HCl, pH 7.4, 150 mM NaCl, 0.2% NP-40) and then re-suspended in 20 μL washing buffer. BODIPY-N3 (1 μL of 2 mM solution in DMF), Tris[(1-benzyl-1H-1,2,3-triazol-4-yl)methyl]amine (1 μL of 10 mM solution in DMF), CuSO_4_ (1 μL of 40 mM solution in H_2_O) and Tris(2-carboxyethyl)phosphine (1 μL of 40 mM solution in H_2_O) were added into the reaction mixture. The click chemistry reaction was allowed to proceed at room temperature for 30 min. Then SDS loading buffer was added and the mixture was heated at 95 °C for 10 min. After centrifugation at 15,000 g for 2 min, the supernatant was collected, treated with 500 mM hydroxylamine, and heated at 95 °C for 5 min. The samples were resolved by 12% SDS-PAGE. BODIPY fluorescence signal was recorded using a Typhoon 9400 Variable Mode Imager (GE Healthcare Life Sciences).

### *In vitro* SHMT2 defatty acylation assay

The SHMT2 plasmid was transfected into HEK293T cells using FuGene 6 transfection reagent. After 24 h, the cells were treated with 50 μM of Alk14 for another 6 h. The cells were harvested and Flag-SHMT2 was affinity purified using anti-Flag affinity gel as described above. After washing the affinity gel three times with washing buffer (25 mM Tris-HCl, pH 7.4, 150 mM NaCl, 0.2% NP-40), 50 μL of triple FLAG peptide solution (300μM in 25 mM Tris-HCl, pH 7.4, 150 mM NaCl) was added to the affinity gel and incubated for 30 min at 4 °C to elute SHMT2 protein. The elution step was repeated two times. 50 μL of purified SHMT2 was incubated with 300 μM of HDAC11 purified from HEK293T cells or buffer for 1 h at 37 °C. After incubation, 200 μL of methanol, 75 μL of chloroform, and 150 μL of water were added to each sample. After vortexing, the samples were centrifuged at 15,000 g for 20 min at 4 °C. The supernatant was gently removed by pipetting, and to each pellet was added 1 mL of methanol. The samples were again vortexed and spun down at 15,000 × g for 10 min at 4 °C. Then methanol was removed and the methanol wash step was repeated. After the second methanol wash, the protein pellets were air-dried for 10-15 min and then re-solubilized in 20 μL of 4% SDS buffer (50 mM triethanolamine at pH 7.4, 150 mM NaCl, 4% (w/v) SDS) and followed by the click chemistry reaction using BODIPY-N3 as mentioned above. The samples were resolved by 12% SDS-PAGE. BODIPY fluorescence signal was recorded using a Typhoon 9400 Variable Mode Imager (GE Healthcare Life Sciences).

### Metabolic Labeling of Mammalian Cells

Cells were treated with 50 μM of Alk14 for 6 h. The cells were collected by centrifugation at 1000 g for 5 min and then lysed in NP-40 lysis buffer (25 mM Tris-HCl, pH 7.4, 150 mM NaCl, 10% glycerol, and 1% NP-40) with protease inhibitor cocktail (1:100 dilution) at 4 °C for 30 min. After centrifuging at 15,000 g for 15 min, the supernatant was collected. The protein concentration was determined using the Pierce BCA Protein Assay Kit. A total of 50 μg of proteins from each sample was aliquoted and adjusted to a final volume of 50 μL with NP-40 lysis buffer. BODIPY-N3 (2.5 μL of 2 mM solution in DMF), Tris[(1-benzyl-1H-1,2,3-triazol-4-yl)methyl]amine (2.5 μL of 10 mM solution in DMF), CuSO_4_ (2.5 μL of 40 mM solution in H_2_O) and Tris(2-carboxyethyl)phosphine (2.5 μL 40 mM solution in H_2_O) were added into the reaction mixture. The click reaction was allowed to proceed at room temperature for 30 min. After incubation, 200 μL of methanol, 75 μL of chloroform, and 150 μL of water were added to each sample. After vortexing, the samples were centrifuged at 15,000 g for 20 min at 4 °C. The supernatant was gently removed by pipetting, and to each pellet was added 1 mL of methanol. The samples were again vortexed and spun down at 15,000 g for 10 min at 4 °C. The methanol was removed, and the protein pellets were washed again with 1 mL of methanol. After the second methanol wash, the protein pellets were air-dried for 15 min and then resolubilized in 50 μL of 4% SDS buffer (50 mM triethanolamine at pH 7.4, 150 mM NaCl, 4% (w/v) SDS). To each sample, 10 μL of 6× loading buffer (374 mM Tris-HCl at pH 6.8, 12% SDS, 600 mM DTT, 60% v/v glycerol, 0.06% bromophenol blue) was added, and the samples were boiled at 95 °C for 5 min. To remove cysteine palmitoylation, 9 μL of each sample was mixed with 1 μL of 5 M hydroxylamine at pH 8.0 and heated at 95 °C for 7 min. The samples were then resolved on 12% SDS-PAGE gel. The gel was destained in 50% (v/v) acetic acid, 40% (v/v) methanol, and 10% (v/v) water for overnight at 4 °C and then destained in water for another 30 min. The fluorescent signal was visualized with a Typhoon 9400 Variable Mode Imager (GE Healthcare Life Sciences).

### Detection of lysine fatty acylation on endogenous SHMT2 by western blot

Cells were treated with or without 50 μM Alk14 for 6 h. The cells were collected by centrifugation at 1000 g for 5 min and then lysed in 4% SDS lysis buffer (50 mM triethanolamine at pH 7.4, 150 mM NaCl, 4% (w/v) SDS) with protease inhibitor cocktail (1:100 dilution) and nuclease (1:1000 dilution) at room temperature for 15 min. The protein concentration was determined by the BCA assay. A total of 1 mg of protein for each sample was used and the volume was brought to 1 ml with 4% SDS lysis buffer. Biotin-N3 (25 μL of 5 mM solution in DMF), Tris[(1-benzyl-1H-1,2,3-triazol-4-yl)methyl]amine (25 μL of 2 mM solution in DMF), CuSO_4_ (25 μL of 50 mM solution in H_2_O) and Tris(2-carboxyethyl)phosphine (12.5 μL of 50 mM solution in H_2_O) were added into the reaction mixture. The click chemistry reaction was allowed to proceed at room temperature for 90 min. After incubation, 5 mL of methanol, 1.875 mL of chloroform, and 3.75 mL of water were added to each sample. After vortexing, the samples were centrifuged at 15,000 g for 20 min at 4 °C. The supernatant was gently removed by pipetting, and to each pellet was added 10 mL of methanol. The samples were again vortexed and spun down at 15,000 g for 10 min at 4 °C. The methanol was removed, and the protein pellets were washed again with 1 mL of methanol. After the second methanol wash, the protein pellets were air-dried for 15 min and then resolubilized in 0.5 mL of 4% SDS buffer. 1 mL of 1% Brij97 buffer (50 mM triethanolamine at pH 7.4, 150 mM NaCl, 1% (w/v) Brij97) was added to dilute the SDS concentration and then mixed with 100 μL of streptavidin agarose beads (ThermoFisher). The mixture was agitated for 1 h at room temperature. After washing the beads three times with washing buffer (PBS with 0.2% SDS), the beads were further treated with 0.4 M hydroxylamine for 1 h at room temperature to remove any cysteine modification. After washing the beads three times with PBS, 20 μL of elution buffer (25 mM Tris-HCl, pH 7.4, 150 mM NaCl, 0.2% NP-40) was added. To each elution, 4 μL of 6× loading buffer was added, and the samples were boiled at 95 °C for 5 min. The samples were further analyzed by western blot against SHMT2.

### SILAC

HDAC11 knockdown or knock out cells were cultured in DMEM with [^13^C_6_,^15^N_2_]-L-lysine and [^13^C_6_,^15^N_4_]-L-arginine for five generations. Wild type cells were cultured in DMEM containing L-lysine and L-arginine for five generations. Then cells were treated with 50 μM Alk14 for 6 h. The cells were collected by centrifugation at 1000 × g for 5 min and then lysed in 4% SDS lysis buffer (50 mM triethanolamine at pH 7.4, 150 mM NaCl, 4% (w/v) SDS) with protease inhibitor cocktail (1:100 dilution) and nuclease (1:1000 dilution) at room temperature for 15 min. The protein concentrations were determined using the Pierce BCA Protein Assay Kit. Then 5 mg of the heavy lysate was mixed with 5 mg of the corresponding light lysate in a 50-ml tube and the volume was brought to 4.45 ml with 4% SDS lysis buffer. Biotin-N3 (100 μL 5 mM solution in DMF), Tris[(1-benzyl-1H-1,2,3-triazol-4-yl)methyl]amine (100 μL of 2 mM solution in DMF), CuSO_4_ (100 μL of 50 mM solution in H_2_O) and Tris(2-carboxyethyl)phosphine (50 μL of 50 mM solution in H_2_O) were added into the reaction mixture. The click chemistry reaction was allowed to proceed at room temperature for 90 min. After incubation, 10 mL of methanol, 3.75 mL of chloroform, and 7.5 mL of water were added to each sample. After vortexing, the samples were centrifuged at 15,000 g for 20 min at 4 °C. The supernatant was gently removed by pipetting, and to each pellet was added 20 mL of methanol. The samples were again vortexed and spun down at 15,000 g for 10 min at 4 °C. The methanol was removed, and the protein pellets were washed again with 1 mL of methanol. After the second methanol wash, the protein pellets were air-dried for 15 min and then resolubilized in 1 mL of 4% SDS buffer. 2 mL of 1% Brij97 buffer (50 mM triethanolamine at pH 7.4, 150 mM NaCl, 1% (w/v) Brij97) was added to dilute the SDS concentration and then mixed with 100 μL streptavidin agarose beads (ThermoFisher). The mixture was rocked for 1 h at room temperature. After washing the beads three times with washing buffer (PBS with 0.2% SDS), the beads were further treated with 0.4 M hydroxylamine for 1 h at room temperature to remove any cysteine modification. After washing beads three times with PBS, 100 μL of elution buffer (25 mM Tris-HCl, pH 7.4, 150 mM NaCl) was added and the samples were boiled at 95 °C for 5 min. The supernatant was obtained and the reduction and alkylation of cysteines were then carried out. Disulfide reduction and protein denaturation were performed in 6 M urea, 10 mM DTT, 50 mM Tris-HCl, pH 8.0, at room temperature for 0.5 h. Then iodoacetamide was added (final concentration, 40 mM) and incubated at room temperature for 0.5 h. After that, 0.5 M of DTT was added to a final concentration of 15 mM and incubated at room temperature for 0.5 h to stop alkylation. The sample was diluted seven times with 50 mM Tris-HCl, pH 8.0, and 1 mM CaCl_2_, before 2 μg trypsin was added and incubated at 37 °C for 18 h. The digestion was quenched with 0.1% trifluoroacetic acid and the mixture was desalted using Sep-Pak C18 cartridge. The lyophilized peptides were used for LC-MS/MS analysis.

### Nano-LC-MS/MS analysis

The lyophilized peptides were dissolved in 2% acetonitrile with 0.5% formic acid for nano-LC-ESI-MS/MS analysis, which was carried out on a LTQ-Orbitrap Elite mass spectrometer (Thermo Fisher Scientific). The Orbitrap was interfaced with Dionex UltiMate3000 MDLC system (Thermo Dionex). Protein samples were injected onto a Acclaim PepMap nano Viper C18 trap column (5 μm, 100 μm × 2 cm, Thermo Dionex) at a flow rate of 20 μL/min for on-line desalting and then separated on C18 RP nano column (5 μm, 75 μm × 50 cm, Magic C18, Bruker). The gradient for HPLC condition was 5−38% acetonitrile with 0.1% formic acid over 120 min. The flow rate was 0.3 μL/min. The Orbitrap Elite was operated in positive ion mode with spray voltage 1.6 kV and source temperature 275 °C. Data-dependent acquisition (DDA) mode was used by one precursor ions MS survey scan from m/z 300 to 1,800 at resolution 60,000 using FT mass analyzer, followed by up to 10 MS/MS scans at resolution 15,000 on 10 most intensive peaks. All data were acquired in Xcalibur 2.2 operation software (Thermo Fisher Scientific).

### SHMT2 enzymatic assay

SHMT2 enzymatic activity was analyzed using a radioactivity assay as reported previously with minor modifications ^37^. Briefly, assay systems contained 10 μM of 3-^14^C-L-Serine (PerkinElmer), 0.25 mM pyridoxal-5-phosphate (Santa Cruz Biotechnology), 2 mM tetrahydrofolate (Cayman), 2 mM Tris(2-carboxyethyl)phosphine and 50 mM potassium phosphate in a total volume of 100 μl, pH 7.4. SHMT2 protein was purified using anti-Flag affinity beads from HEK293T cells. Equal amount of SHMT2 (as analyzed by western blot) attached to beads were incubated at 37 °C for 1 h in the assay system mentioned above. The reaction was terminated with 75 μL of 1 M sodium acetate, 50 μL of 0.1 M formaldehyde, and 75 μL of 0.4 M dimedone. The vessels were heated for 5 min in a boiling water bath to accelerate the formation of the HCHO dimedon derivative. The tubes were then cooled for 5 min in an ice bath before the dimedone compound was extracted by vigorous shaking with 1 mL of toluene. Two minutes of centrifugation separated the phases and 0.8 mL of upper phase were removed for scintillation counting analysis.

### IFNαRI internalization analysis by flow cytometry

Internalization of endogeneous IFNαRI was determined by flow cytometry. 1× 10^5^ Cells were treated with 1000 U/mL IFNα2a for indicated time periods and then the cells were collected immediately on ice. The cells were washed by cold PBS and then blocked by 3% BSA (dissolved in PBS) for 30 min at 4 °C. Cells were then washed and stained with PE conjugated anti-human IFNαRI antibody (PBL assay science) for 45 min at 4 °C. Isotype IgG was used as negative control for flow cytometry analysis. After incubation, the cells were further fixed with 4% paraformaldehyde. All the samples were analyzed using LSRII flow cytometer (BD Biosciences). Flow data was analyzed in FlowJo and relative geometric mean of fluorescent intensity was calculated using following the formula: relative geometric mean of fluorescent intensity (gMFI) = (gMFI of IFNaR antibody stained sample – gMFI of isotype stained sample) / (gMFI of IFNaR antibody stained time 0 sample – gMFI of isotype stained sample).

### IFNαR1 ubiquitination assay

Cells were treated with 1000 U/mL IFNα2a with or without 10 μM cysteine protease inhibitor E64 (Sigma) for 1h. Cells treated with vehicle control were used as negative controls. The cells were collected by centrifugation at 1000 g for 5 min and then lysed in 25μL of 4% SDS lysis buffer (50 mM triethanolamine at pH 7.4, 150 mM NaCl, 4% (w/v) SDS) with protease inhibitor cocktail (1:100 dilution) and nuclease (1:1000 dilution) at room temperature for 15 min. Then 975 μL of NP-40 lysis buffer (25 mM Tris-HCl, pH 7.4, 150 mM NaCl, 10% glycerol, and 1% NP-40) and 20 μL of protein-A/G-agarose beads were added. The sample was incubated in a rotator for 60 min at 4 °C. After centrifuge at 6,000 g for 1 min, the supernatant was collected and quantified the protein concentration using the Pierce BCA Protein Assay Kit. A total of 500 μg of proteins from each sample was aliquoted and adjusted to a final volume of 1000 μL with NP-40 lysis buffer. Anti-human IFNαR1 antibody (LSBio) was added and incubated overnight at 4 °C. Isotype IgG was used as negative control. Then 20 μl of protein-A/G-agrose beads were added the mixture were incubated for 3h at 4 °C. The beads were washed five times with wash buffer (25 mM Tris-HCl, pH 7.4, 150 mM NaCl, 0.2% NP-40) and then re-suspended in 20 μL of wash buffer. To each sample, 4 μL of 6× loading buffer was added, and the samples were boiled at 95 °C for 5 min. The samples were further analyzed by western blot.

### Viral infection assay

Cells were treated with 1000 U/mL IFNα2a for 18h and then washed extensively with fresh media. Lentiviruses containing the pCCL-GFP (multiplicity of infection or MOI = 1, generated from HEK293T) were used to infect cells for 6 h. After removing the virus, cells were then changed to fresh media and incubated for 48 h. The cells were collected by centrifugation at 1000 g for 5 min and then lysed in NP-40 lysis buffer (25 mM Tris-HCl, pH 7.4, 150 mM NaCl, 10% glycerol, and 1% NP-40) with protease inhibitor cocktail (1:100 dilution) at 4 °C for 30 min. After centrifuging at 15,000 g for 15 min, the supernatant was collected. The protein concentration was determined using the BCA assay. A total of 50 μg of proteins from each sample was aliquoted. The expression of GFP protein in cells lysates was analyzed by western blot.

### Expression and purification of human HDAC11 from yeast cells

The human HDAC11 with an added C-terminal His8 tag was cloned into p423 MET25 vector (ATCC, Manassas, VA). The plasmid p423 MET25-HDAC11 was transformed into S. cerevisiae BY4741 strain (OpenBiosystems, Huntsville, AL). The transformed cells were grown in synthetic complete media lacking histidine for 24 hours. Cells were harvested by centrifugation and re-suspended in 20 mM Tris-HCl (pH 8.0), 500 mM NaCl, 10 mM MgCl_2_, 5 mM imidazole, and 0.5 mM phenylmethylsulfonyl fluoride. Cells were lysed using a bead beater (BioSpec Products, Inc., USA), and purified on a BioLogic DuoFlow 10 System (Bio-Rad, Hercules, CA) using a HisTrap HP column (GE Healthcare, Piscataway, NJ) with a linear gradient from 30 mM imidazole to 500 mM imidazole in 30 min. The fractions were collected and dialyzed against 25 mM Tris-HCl buffer (pH 8.0) containing 150 mM NaCl.

### Immunofluorescence and subcellular fractionation

Cells were fixed with 1 ml 4% paraformaldehyde (PFA) at room temperature for 0.5h, followed by washing samples three times in PBS buffer for 5 min each. Block samples in blocking buffer (5% BSA, 0.1% Saponin in PBS) for 0.5h at room temperature. After blocking, diluted primary antibody was applied (primary antibody was diluted in PBS containing 5% BSA and 0.1% Saponin) and the samples were incubated overnight at 4 °C. After washing the samples in PBS with 0.1% Saponin three times for 5 min each, the samples were incubated with fluorochrome-conjugated secondary antibody diluted in 1 mL of PBS with 5% BSA and 0.1% Saponin for another 1 h at room temperature in the dark. The samples were washed five times in PBS with 0.1% Saponin for 5 min each, and then one drop of antifade reagent (with DAPI) was applied. Confocal imaging was performed using a Zeiss LSM710 confocal microscope.

Subcellular fractionation was performed following our previous report with minimal modifications^31^. Briefly, cells were suspended in 500 μL of fractionation buffer (250 mM Sucrose, 20 mM pH 7.4 HEPEs, 10 mM KCl, 2 mM MgCl_2_, 1 mM, EDTA, 1 mM EGTA, and protease inhibitor cocktail) and passed through a 25G needle 20 times using a 1 mL syringe. The samples were placed on ice for 20 min, and then centrifuged at 800 g for 10 min. The Supernatant was transferred into a fresh tube and centrifuged again at 10,000 g for 10 min. The resulting pellet contained the mitochondria and the supernatant was the cytosolic fraction. The pellet was washed with 500 μL of fractionation buffer, and then centrifuged at 10,000 g for 10 min again to isolate the mitochondria. The mitochondria were further lysed in NP-40 lysis buffer (25 mM Tris-HCl, pH 7.4, 150 mM NaCl, 10% glycerol, and 1% NP-40) with protease inhibitor cocktail (1:100 dilution) as mentioned above.

### RNA extraction, reverse transcription and PCR analysis of the mRNA levels of target genes

Total RNAs were extracted using a RNeasy Mini kit (Qiagen). Reverse transcription was performed with SuperScript III First-Strand Synthesis kit (Invitrogen) following the manufacturer’s instructions. For real-time PCR analysis, iTaq Universal SYBR Green Supermix (Bio-Rad) was used following the manufacturer’s instructions. The reaction was monitored using QuantStudio 7 system (Thermofisher). The primers for real-time PCR analysis are listed as follow:

ISG15 forward: CGCAGATCACCCAGAAGATCG

ISG15 reverse: TTCGTCGCATTTGTCCACCA

PKR forward: TGGAAAGCGAACAAGGAGTAAG

PKR reverse: CCAAAGCGTAGAGGTCCACTT

SHMT2 forward: GCCACGGCTCATCATAGCTG

SHMT2 reverse: AGCAGGTGTGCTTTGACTTCA

KIAA0157 forward: CGCAATACGCAGCAGCAGATGTC

KIAA0157 reverse: TGAGTGGAATTGTTGGCAGTGGAG

HDAC11 forward: CACGCTCGCCATCAAGTTTC

HDAC11 reverse: GAAGTCTCGCTCATGCCCATT

GAPDH forward: AATCCCATCACCATCTTCCA

GAPDH reverse: TGGACTCCACGACGTACTCA

Mouse ISG15 forward: GGTGTCCGTGACTAACTCCAT

Mouse ISG15 reverse: TGGAAAGGGTAAGACCGTCCT

Mouse PKR forward: ATGCACGGAGTAGCCATTACG

Mouse PKR reverse: TGACAATCCACCTTGTTTTCGT

Mouse GAPDH forward: AGGTCGGTGTGAACGGATTTG

Mouse GAPDH reverse: TGTAGACCATGTAGTTGAGGTCA

### Viral infection *in vivo.*

C57BL/6 background *Hdac11* constitutive knockout was generated at the Institut Clinique de la Souris (ICS; llkirch, France) in collaboration with Merck & Co. Male and female mice (6-8 weeks of age) were used for the experiments, and wild-type (WT) C57BL/6 mice served as the control. Mice were infected with VSV (Vesicular stomatitis virus, Indiana strain, from ATCC^®^ VR158^TM^, 1*10^8^ PFU per mouse) by intraperitoneal injection. After 36 hours, mice were euthanized and the liver, lung and spleen tissue were collected. All animal experiments were conducted in accordance with the guidelines of the National Advisory Committee on Laboratory Animal Research and the George Washington University Institutional Animal Care and Use Committee.

### Statistical analysis

Data are presented as mean ± s.d. Significance was assessed by two-tailed Student's t-test between two groups; *P < 0.05, **P < 0.01.

## Acknowledgements

This work is support in part by HHMI, Cornell University, and a grant from NIH/NIDDK DK107868. We thank Dr. Sheng Zhang and Dr. Ievgen Motorykin at the Proteomic and MS Facility of Cornell University for help with the SILAC experiments, Cornell University Biotechnology Resource Center (BRC) Imaging Facility for help with the confocal microscopy, which is supported in part by NIH S10RR025502, and Dr. Weishan Huang for help with flow cytometry, and Cornell University NMR Facility, which is supported in part by NSF CHE-1531632. The Orbitrap Fusion mass spectrometer is supported by NIH SIG 1S10 OD017992-01 grant.

## Author contribution

J.C. and H.L. conceived the project. J.C. designed and performed all the biochemical and cellular studies except those noted below. L.S. carried out the virus infection studies in mice. P.A. carried the SILAC experiments. N.A.S synthesized Alk14. X.Z. cloned SHMT2 and helped with Alk14 labeling. H.L. directed and supervised all the biochemical studies. E.S. supervised the mouse studies. J.C and H.L. wrote the manuscript and all authors reviewed, edited, and approved the manuscript.

## Supplementary figures and legends

**Figure S1.**
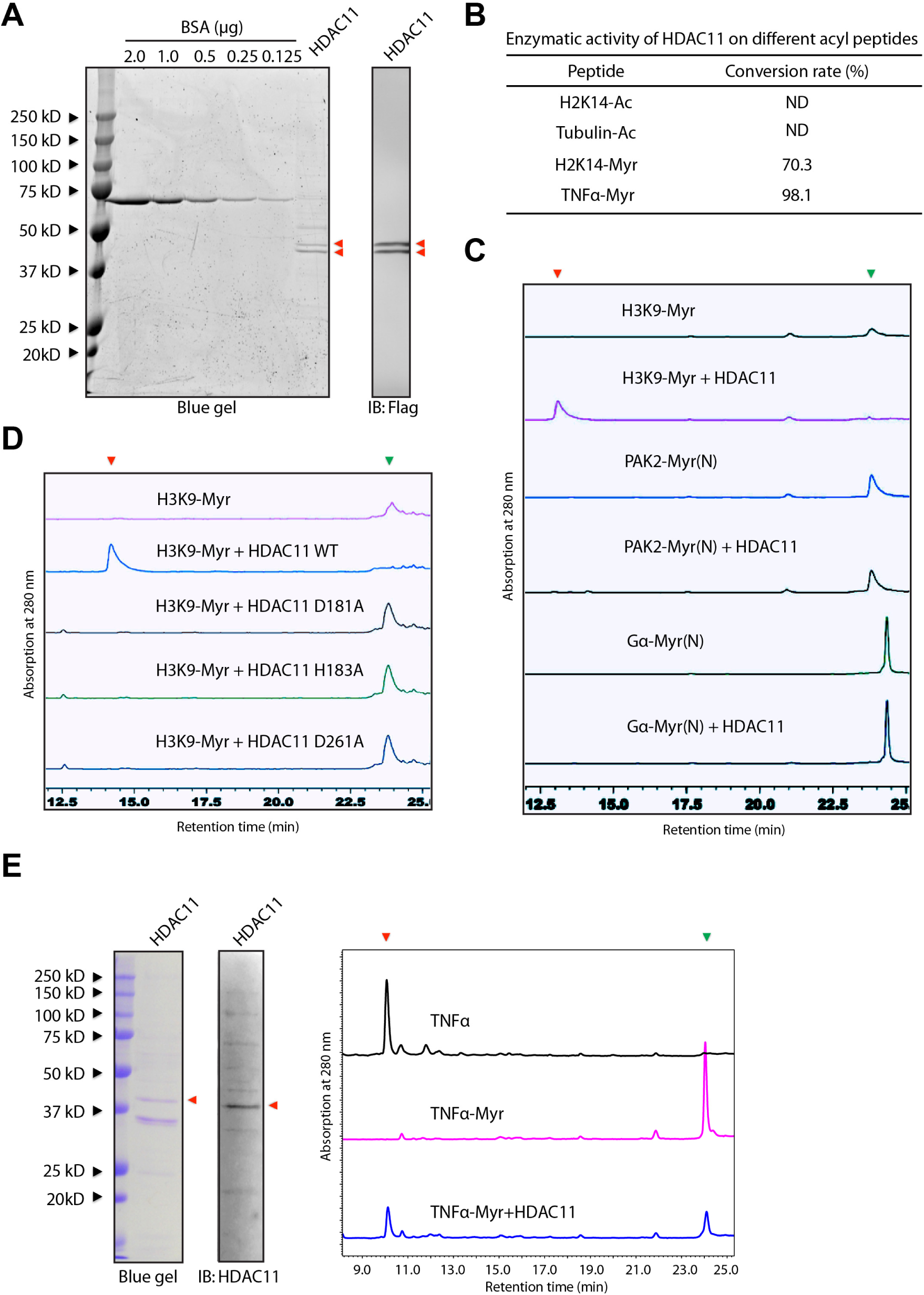
**A.** Flag-tagged HDAC11 purified from 293T cells. *Left,* Coomassie blue stained gel of Flag-tagged HDAC11; *Right,* western blot of Flag-tagged HDAC11 using anti-Flag antibody. **B.** HDAC11 enzymatic activity on different acyl peptides. ND, no detectable activity. **C.** Representative HPLC traces showing the hydrolysis of myristoyl lysine H3K9 peptide, N-myristoyl PAK2 peptide, and N-myristoyl Gα peptide by HDAC11. The experiments were repeated at least twice. **D.** Representative HPLC traces showing the hydrolysis of myristoyl lysine H3K9 peptide by HDAC11 WT and mutants (D181A, H183A, D261A). **E.** The enzymatic activity of His6-tagged HDAC11 purified from yeast on myristoyl lysine TNFα peptide. *Left,* SDS-PAGE gel of His6-tagged HDAC11 and the western blot of His6-tagged HDAC11 using anti-HDAC11 antibody; *Right,* Representative HPLC traces showing the hydrolysis of myristoyl lysine TNFα peptide by HDAC11 purified from yeast.

**Figure S2.**
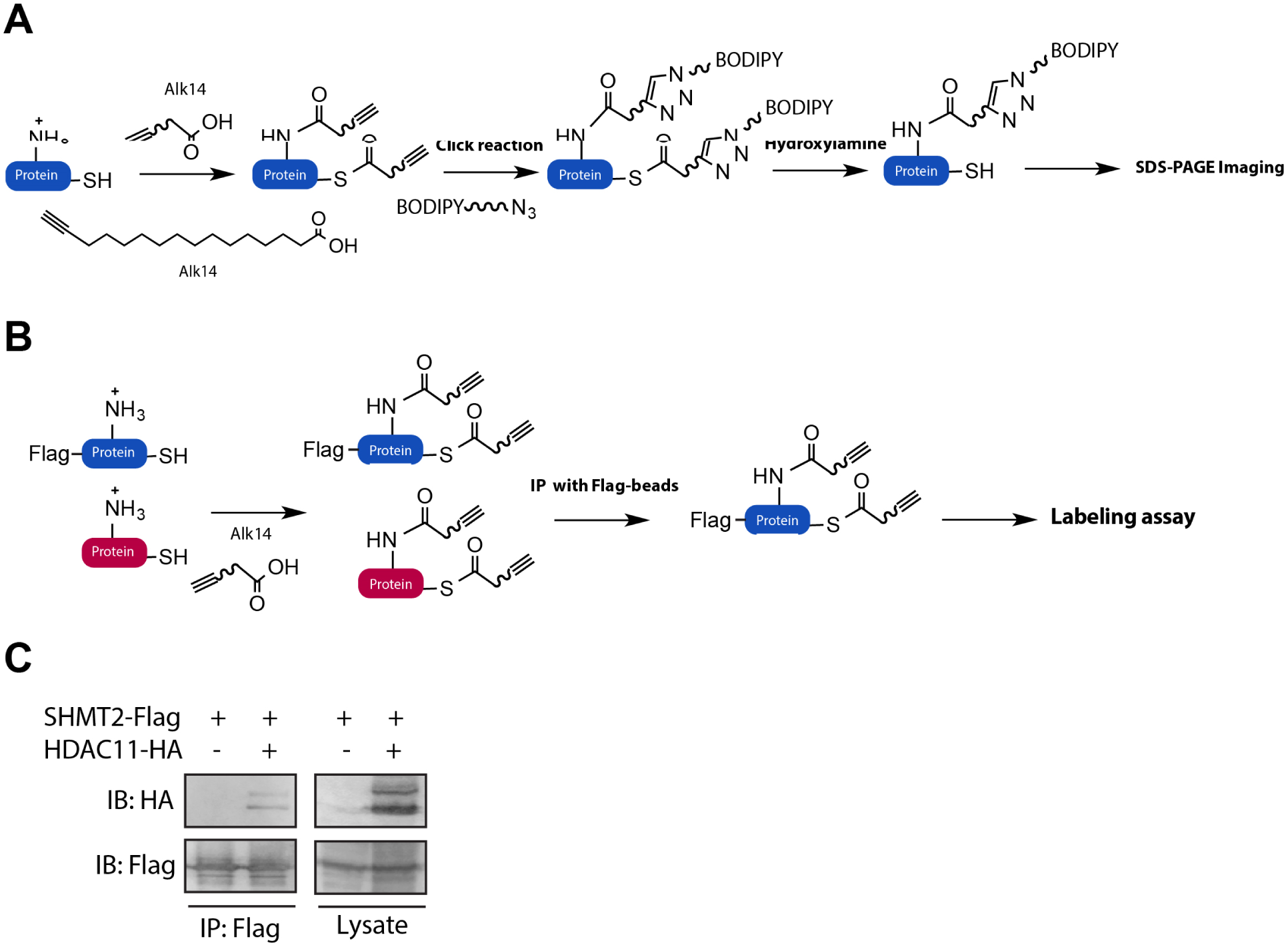
**A.** Scheme showing the metabolic labeling method with the Alk14 probe for in-gel fluorescence analysis of protein lysine fatty-acylation. **B.** The Alk14 probe-based in-gel fluorescence analysis for lysine fatty-acylation of specific protein. **C.** Interaction between ectopic SHMT2 and HDAC11. HEK 293T cells were transfected with Flag-SHMT2 and HA-HDAC11. Cell lysates were immune-precipitated with anti-Flag agarose, followed by immunoblotting with anti-HA and anti-Flag antibodies.

**Figure S3.**
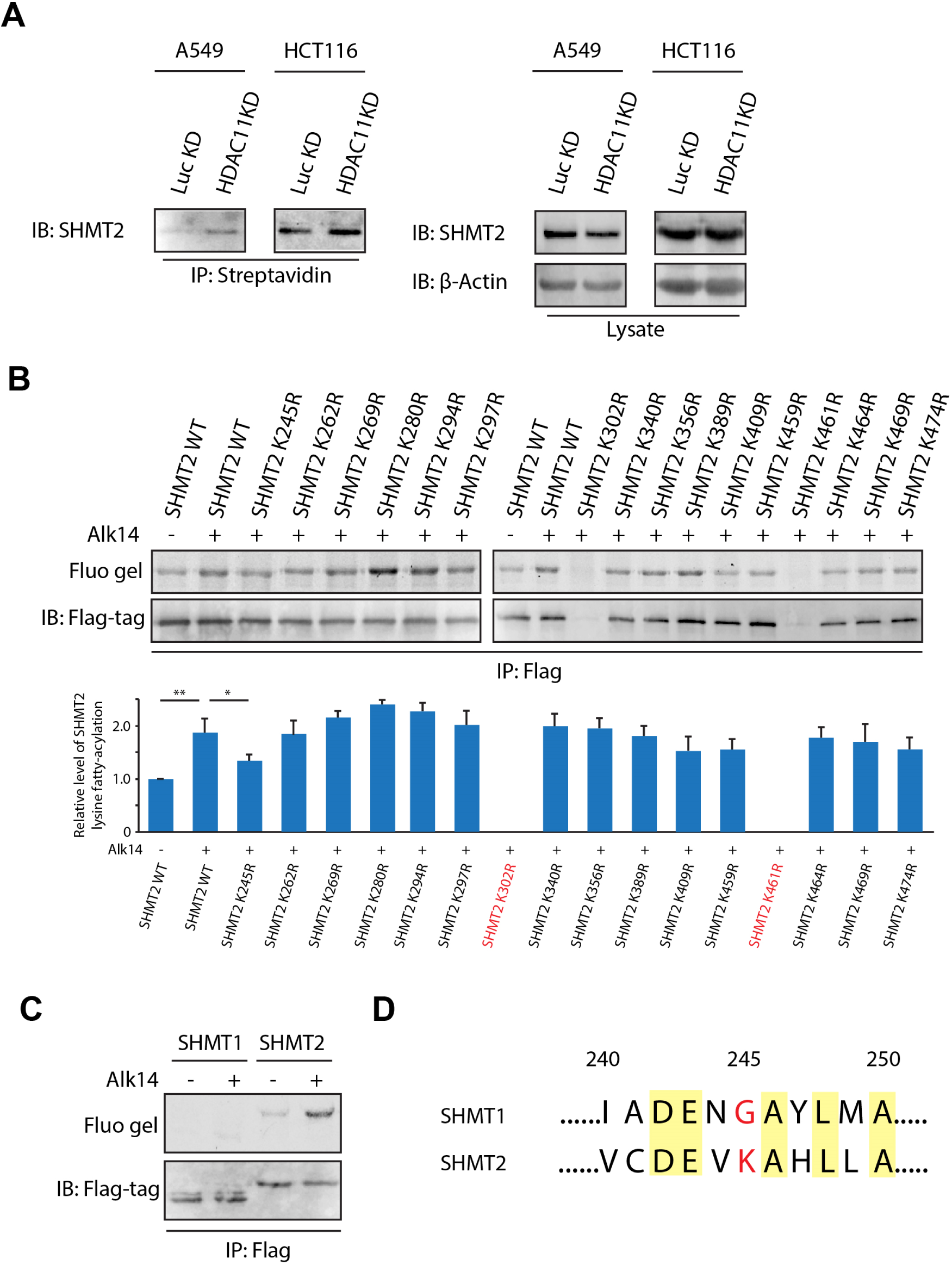
**A.** The fatty-acylation of endogenous SHMT2 was increased after depleting HDAC11 expressions in both A549 and HCT116 cells using Alk14 labeling and biotin pull-down assay as described in methods. **B.** Lysine (K) to Arginine (R) mutation to identify the fatty-acylation site of SHMT2. K245 was the major fatty-acylated site of SHMT2. Quantification of the relative level of lysine fatty-acylation is shown in the right panel. Values with error bars indicate mean ± sd of three replicates. **C.** In-gel fluorescence showing that SHMT2, but not SHMT1, is a lysine fatty-acylated protein. **D.** The sequence alignment of SHMT1 and SHMT2 surrounding K245.

**Figure S4.**
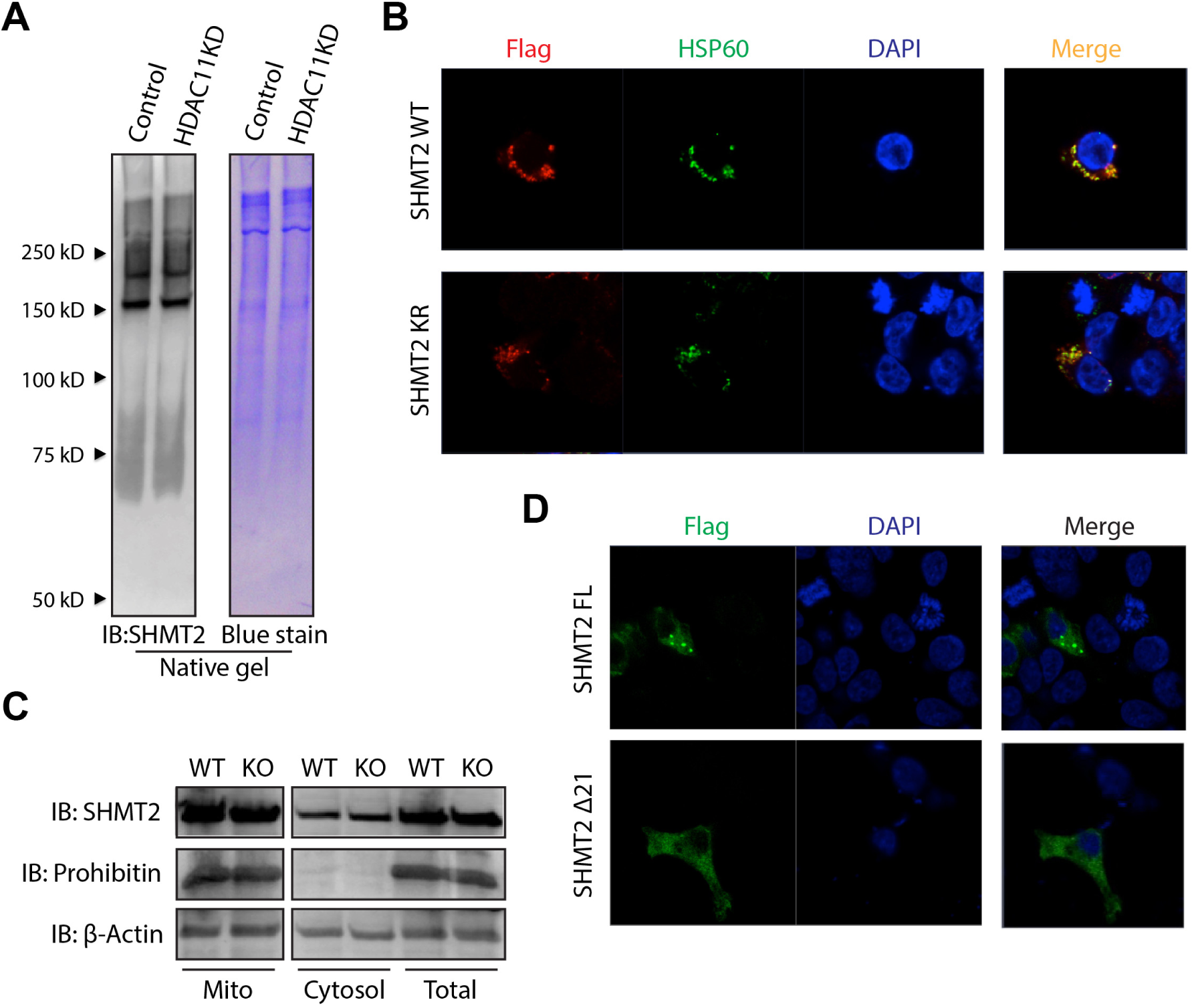
**A.** The oligomeric states of K245R mutant and WT SHMT2 detected using native gel and immunoblotting. B. Colocalization of WT and K245R mutant of SHMT2 with mitochondrial marker protein HSP60. C. Subcellular fractionation followed by immunoblotting using antibody against SHMT2 suggested that a small amount of SHMT2 is localized in the cytosol. D. The Δ21 mutant of SHMT2 is predominantly localized in the cytosol while the full length SHMT2 has cytosolic and strong puncta (mitochondrial) localization.

**Figure S5.**
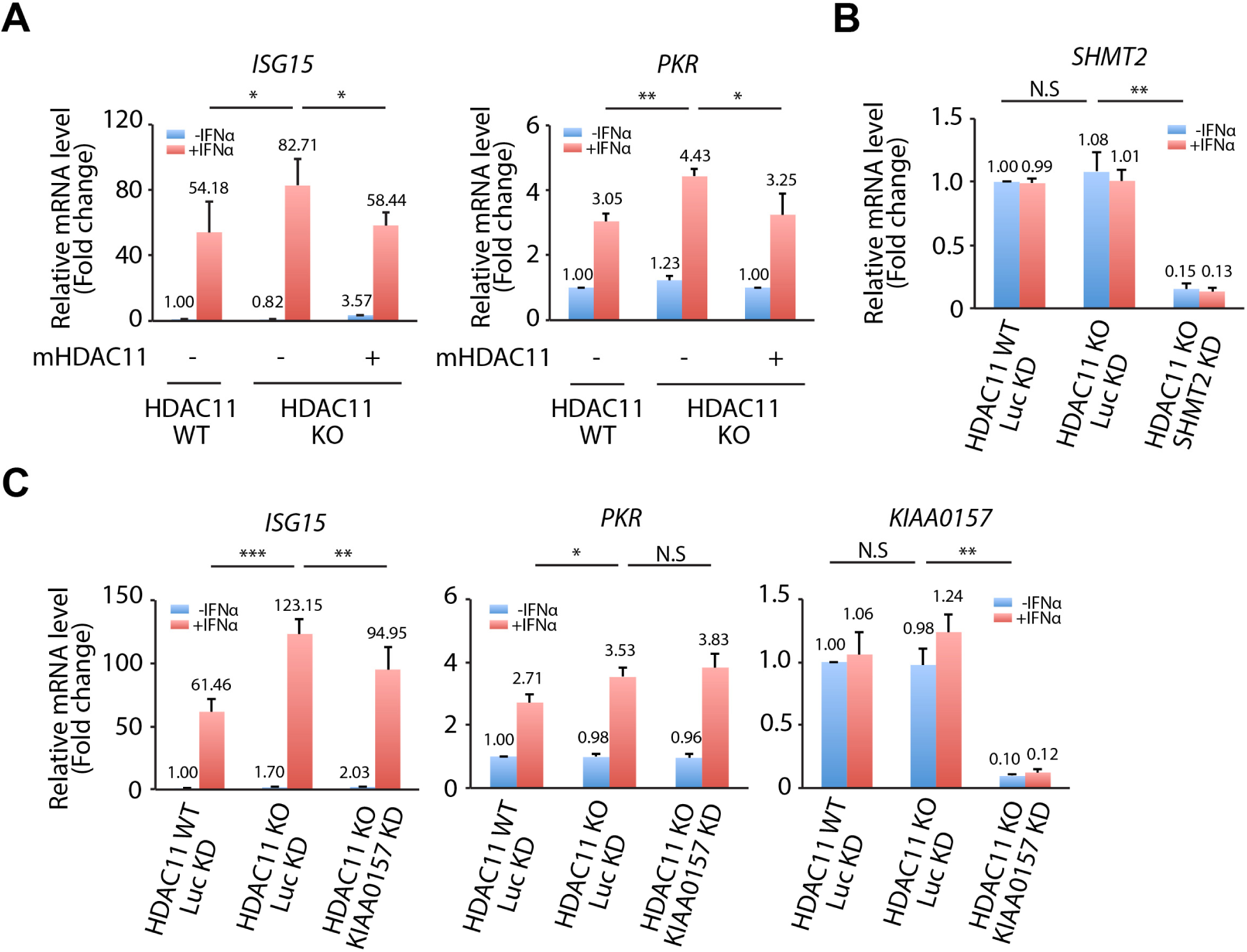
**A.** Re-introducing mouse HDAC11 (mHDAC11) in HDAC11 KO HAP1 cells reverse the increase of *ISG15* and *PKR* mRNA levels. The mRNA levels were quantified by RT-PCR and normalized to GAPDH mRNA levels. B. The knockdown efficiency of ShRNA against SHMT2 in HDAC11 KO HAP1 cells by RT-PCR and normalized to GAPDH mRNA levels. C. The effect of KIAA0157 knockdown on the relative mRNA levels of *ISG15* and *PKR* in HDAC11 KO HAP1 cells. All the experiments were repeated three times. Values with error bars indicate mean ± sd.

**Table S1.** Potential HDAC11 substrates from SILAC1 (HAP1 cells)

**Table S2.** Potential HDAC11 substrates from SILAC2 (HCT116 cells)

**Table S3.** Potential HDAC11 substrates from SILAC3 (MCF-7 cells)

